# The return of the rings: evolutionary role of aromatic residues in liquid-liquid phase separation

**DOI:** 10.1101/2021.10.28.466204

**Authors:** Wen-Lin Ho, Jie-rong Huang

**Affiliations:** Institute of Biochemistry and Molecular Biology, National Yang Ming Chiao Tung University, No. 155 Section 2, Li-nong Street, Taipei, Taiwan; Institute of Biomedical Informatics, National Yang Ming Chiao Tung University, No. 155 Section 2, Li-nong Street, Taipei, Taiwan; Department of Life Sciences and Institute of Genome Sciences, National Yang Ming Chiao Tung University, No. 155 Section 2, Li-nong Street, Taipei, Taiwan

**Keywords:** intrinsically disordered proteins, liquid-liquid phase separation, RNA-binding proteins, biomolecular condensates, membraneless organelle

## Abstract

Aromatic residues appeared relatively late in the evolution of protein sequences. They stabilize the hydrophobic core of globular proteins and are typically absent from intrinsically disordered regions (IDRs). However, recent advances in protein liquid-liquid phase separation (LLPS) studies have shown that aromatic residues in IDRs often act as important “stickers”, promoting multivalent interactions and the formation of higher-order oligomers. To reconcile this apparent contradiction, we compared levels of sequence disorder in RNA binding proteins and the human proteome and found that aromatic residues appear more frequently than expected in the IDRs of RNA binding proteins, which are often found to undergo LLPS. Phylogenetic analysis shows that aromatic residues are highly conserved among chordates, highlighting their importance in LLPS-driven functional assembly. These results suggest therefore that aromatic residues have contributed twice to evolution: in stabilizing structured proteins and in the assembly of biomolecular condensates.

---

Amino acids with aromatic rings stabilize protein structure (Burley and Petsko 1985) and are accordingly rare in intrinsically disordered regions (IDRs)(Dunker, et al. 2001; Yan, et al. 2020). Interestingly however, aromatic residues in IDRs have recently been identified as being crucial in mediating liquid-liquid phase separation (LLPS) (Kwon, et al. 2013; Lin, et al. 2017; Li, Chiang, et al. 2018; Wang, et al. 2018). Here, through protein sequence analyses, we outline how aromatic residues may have contributed twice to evolution.

How did proteins emerge on Earth? Using ferredoxin as an example, Eck and Dayhoff (1996), proposed that early polypeptides were short, with simple compositions, but grew constantly by duplicating their sequences (Eck and Dayhoff 1966). New amino-acids appeared by mutation and the addition of cysteine provided sulfide bonding to ferrous sulfide, a catalyst used as a primitive energy source (Eck and Dayhoff 1966). The more lately incorporated amino acids, including hydrophobic and aromatic residues, enhanced the folding stability of these polypeptides (Burley and Petsko 1985). These polypeptides, or proteins, became the main workhorses of early cells. When lifeforms became more complex with the appearance of eukaryotes and multicellular organisms, proteins became more efficient and “moonlighted” in different functions (Jeffery 1999). The acquisition of IDRs substantially enlarged their functional repertoire (Oldfield and Dunker 2014). The flexibility of IDRs confers many functional advantages to the attached folded domains, notably by increasing the likelihood of interactions with a binding partner (Sugase, et al. 2007) or other molecules (Kriwacki, et al. 1996), in connecting multiple domains (Huang, et al. 2014), and by making the protein more accessible for posttranslational modifications (Li, et al. 2008). Posttranslational modifications alter the physicochemical properties of IDRs (e.g., phosphorylation adds negative charges) and proteins’ propensity to self-assemble (Monahan, et al. 2017; Lu, et al. 2018; Guo, et al. 2019; Saito, et al. 2019). The role of IDRs in tunable and reversible assembly, also known as LLPS, the mechanism that drives the formation of many membraneless organelles (Alberti and Hyman 2021), has only recently been recognized.

The different building blocks of proteins appeared at different stages of evolution. Eck and Dayhoff found for example, that the earliest four amino acids to appear were Ala, Asp, Ser, and Gly (Eck and Dayhoff 1966), and based on multiple criteria, the order of appearance of the amino acids in life is now accepted to be (Trifonov 2004): Gly, Ala, Asp, Val, Pro, Ser, Glu, Leu/Thr, Arg, Ile/Gln/Asn, His, Lys, Cys, Tyr, Met, Trp. Interestingly, as many others have noticed, this suggests that the earliest peptides were disordered, since the most primitive amino acids are not structure-promoting (Zhu, et al. 2016; Kulkarni and Uversky 2018; Katsnelson 2020). These primitive disordered proteins may have initiated life on Earth and acquired aromatic residues later, thereby increasing their folding stability. The initial set of amino acids were implicated again in the evolution of IDRs for various functions. Although aromatic residues appear relatively late in protein evolution, recent studies have shown that they are one of the driving forces of LLPS in the later evolved IDRs (Kwon, et al. 2013; Lin, et al. 2017; Li, Chiang, et al. 2018; Wang, et al. 2018). This apparent contradiction, the appearance of “structure-promoting” amino acids in unstructured regions, raises interesting questions.

We are particularly interested in RNA-binding proteins (RBPs) because evidence is accumulating that the subcellular localization of RNA molecules is mediated by LLPS (Wiedner and Giudice 2021). Posttranscriptional gene control is dependent on RNA molecules, especially mRNA, being in precisely the right place at the right time (Kloc, et al. 2002; Martin and Ephrussi 2009). Accordingly, we analyzed a set of 1,542 RBPs, and a subgroup of 692 mRNA binding proteins (mRBPs) from a census study (Gerstberger, et al. 2014), in comparison with a set of 20,358 human proteins. We collected protein sequences from the UniProt database (UniProt 2019) and analyzed their level of disorder using the PONDR server’s VLXT, VL3, and VSL2 algorithms (Romero, et al. 1997; Radivojac, et al. 2003). We separated residues into ordered and disordered based on the annotation of these algorithms, and calculated the percentage of proteins with sequences of consecutive disordered residues longer than 30, 40, and 50 amino acids (fig. 1*A*; supplementary table S1, Supplementary Material online). Our analysis for the human proteome is similar to Dunker et al.’s classic study of eukaryotic organisms (Dunker, et al. 2000) and is in agreement with the observation that IDRs are more prevalent in RBPs (Varadi, et al. 2015; Zagrovic, et al. 2018). Our results also show that mRBPs have an even higher proportion of disordered residues (fig. 1*A*), with ~10 percentage points (pp) more residues than human proteins in consecutive sequences of 20 or more disordered residues, and ~20 pp more residues found in disordered regions longer than 60 amino acids (supplementary table S1, Supplementary Material online), suggesting that the mRBPs tend to have longer IDRs.

**Fig. 1.**
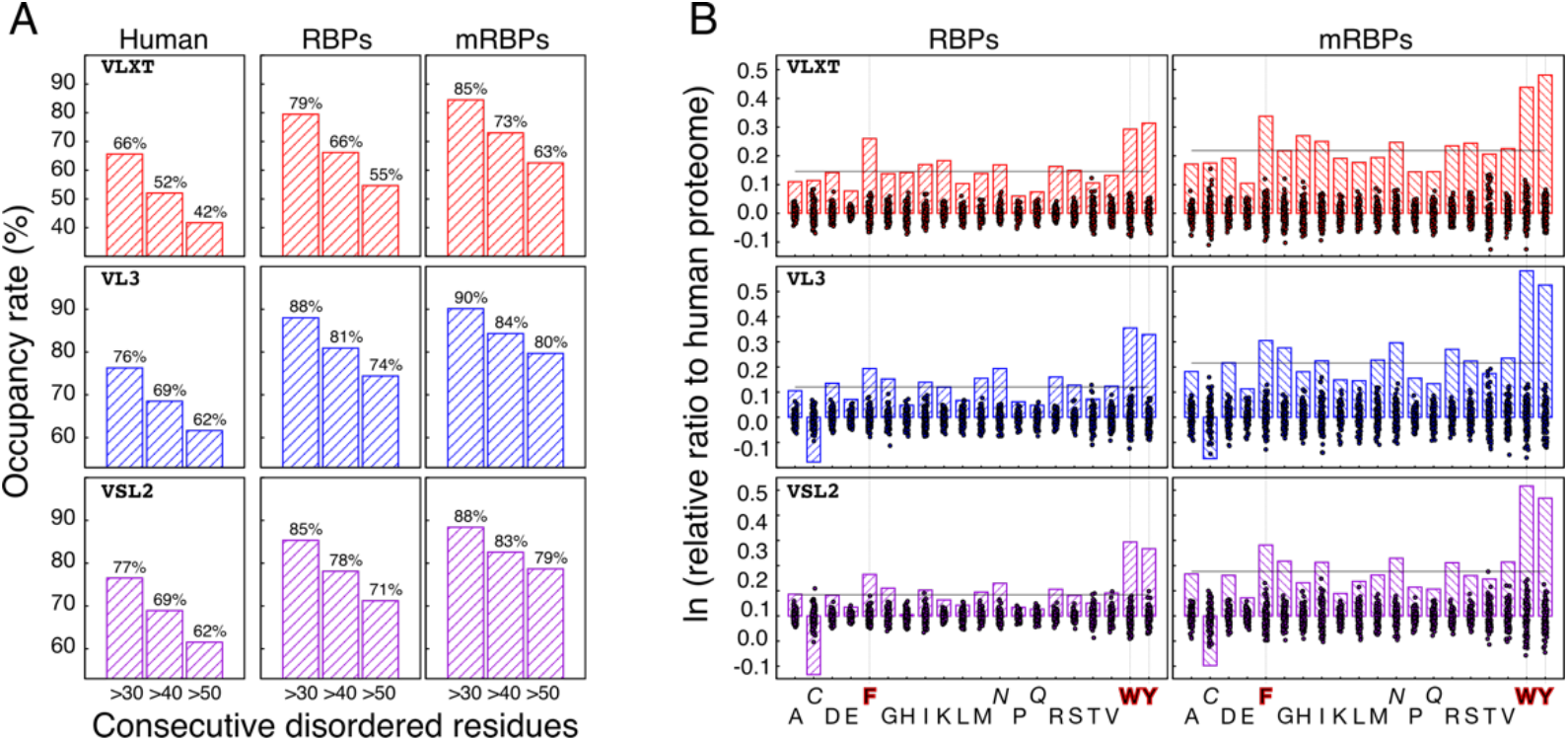
Prevalence of disorder and disorder odds ratios relative to the human proteome by amino acid type for RNA-binding proteins. (A) Proportion of proteins with disordered regions longer than 30, 40, or 50 consecutive residues, as predicted using different algorithms, in the human proteome (left column), RNA-binding proteins (middle column), and mRNA binding proteins (right column). (B) Log-odds ratios relative to the human proteome of being in an IDR for amino acids in RBPs (left) and mRBPs (right). The dashed lines indicate the average value over all amino acid types. The dots represent the values obtained for randomly selected (negative control) subsamples (N = 1542 for RBPs and N = 689 for mRBP, the same numbers as considered in the main analysis) of the human proteome.

Next, we counted the numbers of each amino-acid type in the predicted disordered or folded domains (supplementary tables S2–S4, Supplementary Material online). For each amino acid, we calculated the log-odds ratio relative to the human proteome of occurring in a disordered region in RBPs or mRBPs (fig. 1*B*):

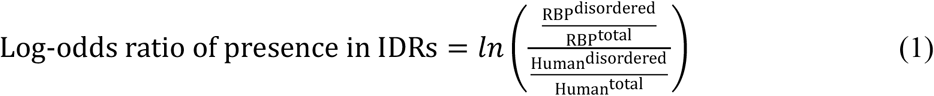

The values obtained are positive for all amino acids (except for cysteine with the VL3 and VSL2 algorithms; fig. 1*B*) and the average values (dashed lines in fig. 1*B*) confirm the above conclusion that IDRs are more prevalent in RBPs than in the human proteome. We also repeated the analysis 1,000 times for random selections of 1,542 or 692 proteins sequences (the numbers of RBPs and mRPBP sequences considered) from the human proteome. The averaged log-odds ratios obtained for the random selections are around zero with standard deviations of mostly less than 0.05 and largest deviations no greater than ± 0.1 (dots in fig. 1*B*; supplementary table S5–S7 and figure S1, Supplementary Material online). This analysis of random selections indicates that the differences for RBPs and mRBPs (bars in fig. 1*B*) are significant.

The results from the different algorithms for each amino acid are consistent, except for cysteine. VL3 and VSL2 predict that the prevalence of cysteines in IDRs is lower in RBPs and mRBPs than in the human proteome, whereas VLXT predicts the opposite, but similar to the range of results of random selection. Although this may simply reflect the algorithm’s use of different scoring functions, it is also possible that IDRs in human proteins have a higher portion of cysteines than those in RBPs do. Note also that asparagine is slightly more likely to be disordered in RBPs and mRBPs, whereas for glutamine the difference is similar to those obtained for the randomly selected pools (fig. 1*B*). Although it has been reported that the IDRs in RBPs are likely to be prion-like, i.e. rich in asparagine and glutamine (Wang, et al. 2018), these two amino acids are not obviously more prevalent than in the human proteome. Among all amino acids, the largest differences obtained with these three algorithms are for phenylalanine, tryptophan, and tyrosine, with values much higher than those obtained for random samples of human proteins, particularly for mRNA targeting proteins (fig. 1*B*). In other words, *structure-promoting* aromatic amino acids are relatively more abundant in *disordered* regions of RBPs than in the human proteome in general.

Functionally important residues are conserved during evolution (Capra and Singh 2007). We therefore aligned orthologue sequences for each RBP and calculated the Shannon sequence entropy of each residue (Sander and Schneider 1991) as a measure of sequence conservation (ranging from 1 for fully conserved to 0 for not conserved at all; fig. 2). The level of conservation is lower in the chordate phylum (dark blue lines in fig. 2) than in mammals (light blue lines). The analysis was repeated (gray lines) for vertebrates (subphylum) and tetrapods (superclass). As expected, folded domains are conserved earlier in evolution than disordered regions as indicated by the dark blue lines being in the regions underlined with a red bar (sequences of more than 40 consecutive disordered residues). We also calculated the average level of conservation for each amino acid. A general increasing trend down the taxonomic ranks is expected. For structured domains, the trends are generally flat, indicating early conservation (examples are shown in figure S2, Supplementary Material online). For disordered regions on the contrary, increases are observed for most amino acids; however, the curves for the aromatic residues in many RBPs are flat, as observed for residues in structured domains (bottom panels in fig 2 and figure S2, Supplementary Material online).

**Fig. 2.**
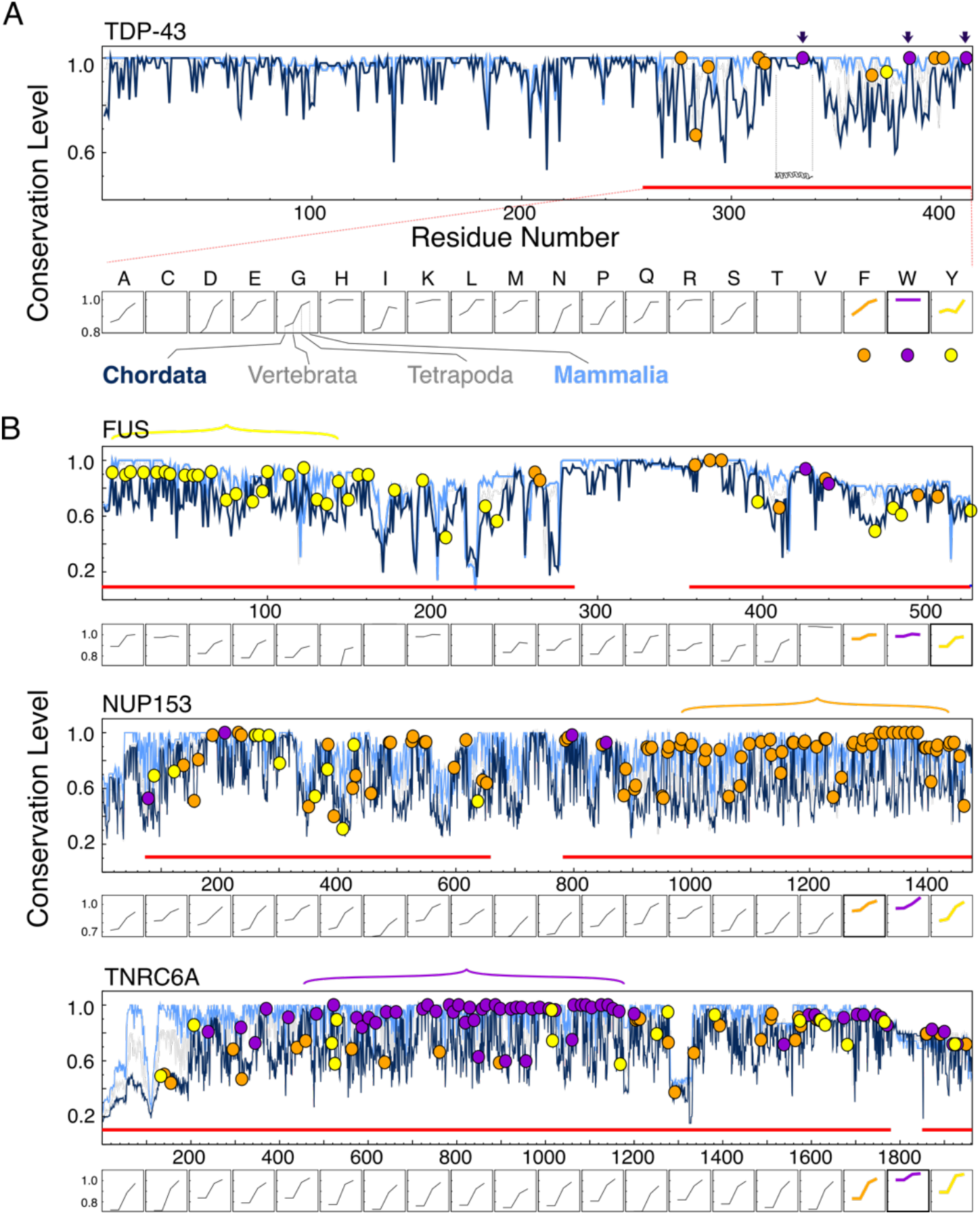
Sequence conservation in example proteins with highly conserved aromatic residues. (A) TDP-43, (B) FUS, NUP153, and TNRC6A. (*Upper panels*) Levels of sequence conservation quantified by the Shannon entropy: a value of 1 means that all the residues aligned at a given position are identical. Levels of conservation in chordates (dark blue), vertebrates (gray), tetrapods (gray), and mammals (light blue), plotted versus the corresponding residue number in the human sequence. Predicted disordered regions longer than 40 residues are indicated with red bars. Aromatic residues are labeled on the chordate line: Phe (orange), Trp (purple), Tyr (yellow). (A) The three arrows indicate tryptophans experimentally identified as being crucial to liquid-liquid phase separation; the transient α-helical region that also contributes to self-assembly is also labeled. (B) The colored horizontal curly brackets indicate highly conserved aromatic-residue-rich regions. (*Lower panels*) Average levels of conservation as a function of decreasing taxonomic rank for amino-acid types in regions predicted to be disordered (indicated by red bars in the upper panel).

In TDP-43 for example, one of the most extensively studied RBP undergoing LLPS (Conicella, et al. 2016; McGurk, et al. 2018; Babinchak, et al. 2019; Sun, et al. 2019), the aromatic residues in its IDRs are highly conserved (fig. 2*A*; the orange/purple/yellow circles on the Chordata line respectively indicate Phe/Trp/Tyr). The three tryptophans known to be key residues in driving LLPS (Li, Chiang, et al. 2018) are conserved in all chordates. The phenylalanines are conserved early but to a lesser extent, in keeping with the fact that they contribute less to LLPS (Li, Chiang, et al. 2018). This interpretation that sequence conservation reflects functional importance is further supported by the fact that the transient α-helical region in TDP-43’s IDR (fig. 2*A*, ~residues 320-340), which is involved in LLPS (Chen, et al. 2016; Conicella, et al. 2016; Li, Chen, et al. 2018) is also conserved.

Many other RBPs have highly conserved aromatic residues, whose functional importance has been demonstrated in several cases (fig. 2*B*). Tyrosines are known to be involved in the well-studied LLPS of the N-terminal domain of the RBP FUS (Lin, et al. 2017; Qamar, et al. 2018; Wang, et al. 2018) and a recent analysis of mammalian FUS proteins found the same trend as observed here (Dasmeh and Wagner 2021). Phe-Gly repeats are a common feature of nucleoporins, which regulate nucleocytoplasmic transport in the nuclear pore complex (Milles, et al. 2015; Onischenko, et al. 2017; Hayama, et al. 2018). The nucleoporin NUP153 is also categorized as an RBP because its Phe-Gly repeats mediate mRNA trafficking (Bastos, et al. 1996). Phenylalanines are conserved in NUP153 (fig. 2*B*) and it is reasonable to suppose that this is also the case in other nucleoporins. The tryptophan-rich region in TNRC6 interacts with Ago2 to promote the phase separation of the miRNA induced silencing complex (Sheu-Gruttadauria and MacRae 2018); all but a few of these tryptophans are highly conserved in chordates (fig. 2*B*). Other examples include the LLPS of CPEB (Ford, et al. 2019) and HNRNPD (Batlle, et al. 2020). Although the role of aromatic residues has not been explicitly studied in these proteins, their conservation in chordates hints at a possible role in LLPS related functions (figure S3, Supplementary Material online). Many RBPs that have not been reported to undergo LLPS also have conserved aromatic patterns, including RBM19, DDX18, KHDRBS1, ABT1, RSL24D1, SMAD5, and TAF9 (figure S2*B* and S3, Supplementary Material online). We suggest these conserved aromatic residues in IDRs may be involved in LLPS-related functions.

Intrinsically disordered regions appeared late in proteins and RBPs evolved under the selective pressure to assemble precisely in specific cellular locations. This can be achieved by mimicking prion properties (Patel, et al. 2015) or by having short α-helical motifs (Conicella, et al. 2016), blocked charged-pattern (Greig, et al. 2020), or coiled-coil domains (Fang, et al. 2019), features that provide multivalency and promote the formation of higher-order oligomers. Aromatic residues, with weak *π*-*π* or cation-*π* interactions, appear to have been selected in this role, affording the weak, reversible interactions required for biomolecular condensate formation. After contributing a first time in the evolution of folded proteins, the subsequent return of the (aromatic) rings in RBPs appears to have been crucial to the emergence of liquid-liquid phase separation.

## Methods

All human protein sequences were retrieved from UniProt (UniProtKB_2021_01, download date: 2021/02/06, 20396 sequences in total). Disorder predictions were performed using the VLXT, VL3, and VSL2 algorithms on the PONDR web server (Romero, et al. 1997; Radivojac, et al. 2003), and analyzed using in-house scripts.

Human RBP orthologues were obtained from the Orthologous Matrix database (Altenhoff, et al. 2021) using the human protein’s UniProt ID as input. The orthologues were filtered by taxonomy. The sequences of the orthologues were aligned using the Clustal Omega (v.1.2.4) (Sievers, et al. 2011) module in Biopython (Cock, et al. 2009). The aligned sequences were mapped to the human sequence’s residue number and the level of conservation of each residue was quantified by its Shannon entropy (Sander and Schneider 1991).

## Supplementary Material

Supplementary data are available at Molecular Biology and Evolution online.

## Data Availability

All scripts and data generated or analyzed is this study are available in the repository: http://github.com/allmwh/the_return_of_the_rings

## Acknowledgements

This research was funded by the Ministry of Science and Technology of Taiwan, grant numbers 109-2113-M-010-003 and 110-2113-M-A49A-504-MY3.

