## Supporting Information for "The return of the rings: evolutionary role of aromatic residues in liquid-liquid phase separation"

**Table S1.** Proportions of proteins with predicted IDRs longer than certain lengths.

| Algorithm | Length longer than | Human proteins (%) | RNA binding proteins (%) | (RBP – Human) (%) | mRNA binding proteins (%) | (mRBP – Human) (%) |
|---|---|---|---|---|---|---|
| VLXT | **20** | **81.8** | **92.6** | 10.9 | **93.6** | 11.8 |
| | **30** | **65.6** | **79.5** | 13.9 | **84.5** | 18.9 |
| | **40** | **52.0** | **66.2** | 14.2 | **73.0** | 21.0 |
| | **50** | **41.7** | **54.7** | 12.9 | **62.6** | 20.8 |
| | **60** | **33.7** | **45.9** | 12.2 | **55.4** | 21.7 |
| VL3 | **20** | **83.3** | **93.4** | 10.1 | **93.9** | 10.6 |
| | **30** | **76.3** | **88.0** | 11.7 | **90.1** | 13.8 |
| | **40** | **68.5** | **80.9** | 12.4 | **84.3** | 15.8 |
| | **50** | **61.6** | **74.4** | 12.8 | **79.7** | 18.1 |
| | **60** | **55.7** | **66.6** | 10.9 | **74.2** | 18.5 |
| VSL2 | **20** | **84.1** | **91.8** | 7.7 | **93.9** | 9.8 |
| | **30** | **76.5** | **85.3** | 8.8 | **88.4** | 11.9 |
| | **40** | **68.9** | **78.1** | 9.2 | **82.6** | 13.7 |
| | **50** | **61.5** | **71.2** | 9.7 | **78.7** | 17.1 |
| | **60** | **55.6** | **64.3** | 8.7 | **73.4** | 17.7 |

**Table S2.** Number of residues predicted to be in ordered/disordered regions by the VLXT algorithm

| | RBPs | | mRBPs | | Human proteins | |
|---|---|---|---|---|---|---|
| | Ordered | Disordered | Ordered | Disordered | Ordered | Disordered |
| A | 33767 | 27760 | 15945 | 14705 | 474415 | 321897 |
| C | 12353 | 2662 | 5358 | 1244 | 220009 | 41333 |
| D | 27224 | 18481 | 13262 | 9798 | 349268 | 188768 |
| E | 31856 | 36215 | 15343 | 18489 | 409580 | 396968 |
| F | 25340 | 4883 | 12221 | 2586 | 362796 | 51625 |
| G | 32066 | 26379 | 15778 | 15089 | 452912 | 293467 |
| H | 15438 | 6544 | 6995 | 3577 | 220858 | 76954 |
| I | 26308 | 11070 | 12352 | 5838 | 369522 | 123116 |
| K | 34806 | 26359 | 16136 | 12385 | 417435 | 233546 |
| L | 54425 | 25927 | 24430 | 12995 | 802230 | 329184 |
| M | 10887 | 8169 | 5332 | 4419 | 151790 | 90374 |
| N | 21112 | 11254 | 10520 | 6336 | 287931 | 119746 |
| P | 25046 | 31437 | 11796 | 18099 | 341273 | 375933 |
| Q | 25311 | 18377 | 12474 | 10261 | 330112 | 211467 |
| R | 26230 | 30610 | 12211 | 16772 | 347223 | 292901 |
| S | 34805 | 36396 | 16745 | 21500 | 529363 | 416555 |
| T | 26013 | 17415 | 12199 | 9710 | 388457 | 219237 |
| V | 34359 | 17768 | 15600 | 9331 | 474983 | 202342 |
| W | 6853 | 2006 | 2992 | 1062 | 114749 | 23332 |
| Y | 18116 | 5406 | 8934 | 3333 | 251740 | 50805 |

**Table S3.** Number of residues predicted to be in ordered/disordered regions by the VL3 algorithm

| | RBPs | | mRBPs | | Human proteins | |
|---|---|---|---|---|---|---|
| | Ordered | Disordered | Ordered | Disordered | Ordered | Disordered |
| A | 34295 | 27232 | 16011 | 14639 | 479203 | 317109 |
| C | 10891 | 4124 | 4765 | 1837 | 175643 | 85699 |
| D | 25476 | 20229 | 11988 | 11072 | 330080 | 207956 |
| E | 31199 | 36872 | 14727 | 19105 | 399940 | 406608 |
| F | 22220 | 8003 | 10423 | 4384 | 324013 | 90408 |
| G | 29712 | 28733 | 13694 | 17173 | 431350 | 315029 |
| H | 13748 | 8234 | 6048 | 4524 | 191475 | 106337 |
| I | 27452 | 9926 | 12931 | 5259 | 378838 | 113800 |
| K | 30335 | 30830 | 13731 | 14790 | 360043 | 290938 |
| L | 53010 | 27342 | 23653 | 13772 | 771505 | 359909 |
| M | 11504 | 7552 | 5595 | 4156 | 159982 | 82182 |
| N | 19550 | 12816 | 9461 | 7395 | 274701 | 132976 |
| P | 23244 | 33239 | 10572 | 19323 | 320476 | 396730 |
| Q | 22870 | 20818 | 10930 | 11805 | 295531 | 246048 |
| R | 27151 | 29689 | 12085 | 16898 | 355342 | 284782 |
| S | 31099 | 40102 | 14545 | 23700 | 477217 | 468701 |
| T | 25201 | 18227 | 11733 | 10176 | 370569 | 237125 |
| V | 34809 | 17318 | 15676 | 9255 | 478562 | 198763 |
| W | 6619 | 2240 | 2768 | 1286 | 113612 | 24469 |
| Y | 16578 | 6944 | 7858 | 4409 | 238289 | 64256 |

**Table S4.** Number of residues predicted to be in ordered/disordered regions by the VLS2 algorithm

| | RBPs | | mRBPs | | Human proteins | |
|---|---|---|---|---|---|---|
| | Ordered | Disordered | Ordered | Disordered | Ordered | Disordered |
| A | 30642 | 30885 | 13975 | 16675 | 429548 | 366764 |
| C | 11246 | 3769 | 4884 | 1718 | 178491 | 82851 |
| D | 22064 | 23641 | 10146 | 12914 | 281602 | 256434 |
| E | 26089 | 41982 | 12175 | 21657 | 326165 | 480383 |
| F | 21578 | 8645 | 10046 | 4761 | 313891 | 100530 |
| G | 25897 | 32548 | 11723 | 19144 | 374088 | 372291 |
| H | 12557 | 9425 | 5427 | 5145 | 170785 | 127027 |
| I | 27085 | 10293 | 12597 | 5593 | 370211 | 122427 |
| K | 25112 | 36053 | 11275 | 17246 | 290910 | 360071 |
| L | 52104 | 28248 | 22962 | 14463 | 750188 | 381226 |
| M | 10076 | 8980 | 4834 | 4917 | 138373 | 103791 |
| N | 16953 | 15413 | 7988 | 8868 | 237124 | 170553 |
| P | 19526 | 36957 | 8703 | 21192 | 263731 | 453475 |
| Q | 19952 | 23736 | 9358 | 13377 | 255138 | 286441 |
| R | 24648 | 32192 | 10737 | 18246 | 314028 | 326096 |
| S | 24465 | 46736 | 11073 | 27172 | 373370 | 572548 |
| T | 22428 | 21000 | 10245 | 11664 | 328426 | 279268 |
| V | 34945 | 17182 | 15635 | 9296 | 473431 | 203894 |
| W | 6355 | 2504 | 2622 | 1432 | 108987 | 29094 |
| Y | 15972 | 7550 | 7451 | 4816 | 228179 | 74366 |

**Table S5**. Summary statistics for the log-odds ratios relative to the human proteome of being in a disordered region as predicted by the VXLT algorithm for randomly sampled subgroups of the human proteome

| | Random sample size = 1542 | | | | Random sample size =689 | | | |
|---|---|---|---|---|---|---|---|---|
| | mean | std | max | min | mean | std | max | min |
| A | 0.00 | 0.02 | 0.05 | -0.05 | 0.00 | 0.03 | 0.08 | -0.09 |
| C | 0.00 | 0.03 | 0.10 | -0.09 | 0.00 | 0.05 | 0.15 | -0.15 |
| D | 0.00 | 0.02 | 0.08 | -0.06 | 0.00 | 0.03 | 0.09 | -0.11 |
| E | 0.00 | 0.01 | 0.04 | -0.05 | 0.00 | 0.02 | 0.07 | -0.08 |
| F | 0.00 | 0.03 | 0.08 | -0.08 | 0.00 | 0.04 | 0.12 | -0.13 |
| G | 0.00 | 0.02 | 0.08 | -0.07 | 0.00 | 0.04 | 0.13 | -0.12 |
| H | 0.00 | 0.03 | 0.08 | -0.08 | 0.00 | 0.04 | 0.18 | -0.13 |
| I | 0.00 | 0.02 | 0.08 | -0.06 | 0.00 | 0.04 | 0.13 | -0.10 |
| K | 0.00 | 0.02 | 0.06 | -0.06 | 0.00 | 0.03 | 0.09 | -0.10 |
| L | 0.00 | 0.02 | 0.06 | -0.07 | 0.00 | 0.03 | 0.10 | -0.11 |
| M | 0.00 | 0.02 | 0.08 | -0.06 | 0.00 | 0.03 | 0.11 | -0.08 |
| N | 0.00 | 0.02 | 0.07 | -0.07 | 0.00 | 0.03 | 0.11 | -0.13 |
| P | 0.00 | 0.02 | 0.06 | -0.05 | 0.00 | 0.03 | 0.10 | -0.09 |
| Q | 0.00 | 0.02 | 0.07 | -0.06 | 0.00 | 0.03 | 0.09 | -0.12 |
| R | 0.00 | 0.02 | 0.05 | -0.06 | 0.00 | 0.02 | 0.09 | -0.08 |
| S | 0.00 | 0.02 | 0.07 | -0.06 | 0.00 | 0.03 | 0.11 | -0.11 |
| T | 0.00 | 0.04 | 0.16 | -0.09 | 0.00 | 0.06 | 0.22 | -0.14 |
| V | 0.00 | 0.02 | 0.06 | -0.07 | 0.00 | 0.03 | 0.09 | -0.11 |
| W | 0.00 | 0.03 | 0.12 | -0.13 | 0.00 | 0.05 | 0.15 | -0.13 |
| Y | 0.00 | 0.03 | 0.09 | -0.08 | 0.00 | 0.04 | 0.13 | -0.12 |

The values are derived from 1000 randomly selected subgroups of the human proteome with sample sizes of 1542 or 689 (the same as those of the samples of RBPs and mRBPs considered) using eq. (1). The mean, standard deviation (std), maximum (max), and minimum (min) values are listed.

**Table S6**. Summary statistics for the log-odds ratios relative to the human proteome of being in a disordered region as predicted by the VL3 algorithm for randomly sampled subgroups of the human proteome

| | Random sample size = 1542 | | | | Random sample size =689 | | | |
|---|---|---|---|---|---|---|---|---|
| | mean | std | max | min | mean | std | max | min |
| A | 0.00 | 0.03 | 0.08 | -0.08 | 0.00 | 0.04 | 0.11 | -0.13 |
| C | 0.00 | 0.04 | 0.11 | -0.10 | 0.00 | 0.06 | 0.17 | -0.19 |
| D | 0.00 | 0.03 | 0.10 | -0.08 | 0.00 | 0.04 | 0.11 | -0.17 |
| E | 0.00 | 0.02 | 0.06 | -0.07 | 0.00 | 0.03 | 0.10 | -0.11 |
| F | 0.00 | 0.04 | 0.14 | -0.12 | 0.00 | 0.05 | 0.14 | -0.17 |
| G | 0.00 | 0.03 | 0.12 | -0.11 | 0.00 | 0.04 | 0.10 | -0.16 |
| H | 0.00 | 0.03 | 0.10 | -0.10 | 0.00 | 0.05 | 0.13 | -0.19 |
| I | 0.00 | 0.04 | 0.14 | -0.12 | 0.00 | 0.06 | 0.17 | -0.19 |
| K | 0.00 | 0.03 | 0.08 | -0.07 | 0.00 | 0.04 | 0.11 | -0.14 |
| L | 0.00 | 0.03 | 0.10 | -0.08 | 0.00 | 0.05 | 0.14 | -0.14 |
| M | 0.00 | 0.03 | 0.09 | -0.09 | 0.00 | 0.05 | 0.14 | -0.17 |
| N | 0.00 | 0.03 | 0.09 | -0.09 | 0.00 | 0.05 | 0.14 | -0.18 |
| P | 0.00 | 0.02 | 0.06 | -0.06 | 0.00 | 0.03 | 0.09 | -0.11 |
| Q | 0.00 | 0.02 | 0.08 | -0.07 | 0.00 | 0.04 | 0.13 | -0.13 |
| R | 0.00 | 0.02 | 0.07 | -0.09 | 0.00 | 0.04 | 0.10 | -0.13 |
| S | 0.00 | 0.03 | 0.08 | -0.07 | 0.00 | 0.04 | 0.12 | -0.13 |
| T | 0.00 | 0.04 | 0.14 | -0.11 | -0.01 | 0.06 | 0.21 | -0.19 |
| V | 0.00 | 0.03 | 0.12 | -0.10 | 0.00 | 0.05 | 0.13 | -0.17 |
| W | 0.00 | 0.04 | 0.16 | -0.12 | 0.00 | 0.07 | 0.20 | -0.23 |
| Y | 0.00 | 0.04 | 0.12 | -0.12 | 0.00 | 0.06 | 0.16 | -0.21 |

The values are derived from 1000 randomly selected subgroups of the human proteome with sample sizes of 1542 or 689 (the same as those of the samples of RBPs and mRBPs considered) using eq. (1). The mean, standard deviation (std), maximum (max), and minimum (min) values are listed.

**Table S7**. Summary statistics for the log-odds ratios relative to the human proteome of being in a disordered region as predicted by the VSL2 algorithm for randomly sampled subgroups of the human proteome

| | Random sample size = 1542 | | | | Random sample size =689 | | | |
|---|---|---|---|---|---|---|---|---|
| | mean | std | max | min | mean | std | max | min |
| A | 0.00 | 0.02 | 0.06 | -0.07 | 0.00 | 0.03 | 0.11 | -0.09 |
| C | 0.00 | 0.04 | 0.14 | -0.13 | 0.00 | 0.06 | 0.14 | -0.21 |
| D | 0.00 | 0.02 | 0.07 | -0.05 | 0.00 | 0.03 | 0.12 | -0.10 |
| E | 0.00 | 0.01 | 0.05 | -0.05 | 0.00 | 0.02 | 0.07 | -0.08 |
| F | 0.00 | 0.03 | 0.10 | -0.11 | 0.00 | 0.05 | 0.15 | -0.14 |
| G | 0.00 | 0.02 | 0.07 | -0.07 | 0.00 | 0.03 | 0.12 | -0.10 |
| H | 0.00 | 0.02 | 0.08 | -0.07 | 0.00 | 0.04 | 0.12 | -0.11 |
| I | 0.00 | 0.03 | 0.10 | -0.13 | 0.00 | 0.05 | 0.16 | -0.16 |
| K | 0.00 | 0.02 | 0.05 | -0.05 | 0.00 | 0.03 | 0.10 | -0.07 |
| L | 0.00 | 0.03 | 0.09 | -0.08 | 0.00 | 0.04 | 0.15 | -0.12 |
| M | 0.00 | 0.02 | 0.07 | -0.07 | 0.00 | 0.03 | 0.11 | -0.10 |
| N | 0.00 | 0.02 | 0.06 | -0.08 | 0.00 | 0.03 | 0.10 | -0.11 |
| P | 0.00 | 0.02 | 0.05 | -0.05 | 0.00 | 0.02 | 0.09 | -0.08 |
| Q | 0.00 | 0.02 | 0.07 | -0.07 | 0.00 | 0.03 | 0.10 | -0.09 |
| R | 0.00 | 0.02 | 0.07 | -0.05 | 0.00 | 0.03 | 0.08 | -0.10 |
| S | 0.00 | 0.02 | 0.05 | -0.05 | 0.00 | 0.03 | 0.09 | -0.08 |
| T | 0.00 | 0.03 | 0.11 | -0.09 | 0.00 | 0.05 | 0.18 | -0.14 |
| V | 0.00 | 0.03 | 0.10 | -0.10 | 0.00 | 0.05 | 0.18 | -0.13 |
| W | 0.00 | 0.04 | 0.12 | -0.12 | 0.00 | 0.06 | 0.17 | -0.16 |
| Y | 0.00 | 0.03 | 0.12 | -0.10 | 0.00 | 0.05 | 0.15 | -0.16 |

The values are derived from 1000 randomly selected subgroups of the human proteome with sample sizes of 1542 or 689 (the same as those of the samples of RBPs and mRBPs considered) using eq. (1). The mean, standard deviation (std), maximum (max), and minimum (min) values are listed.

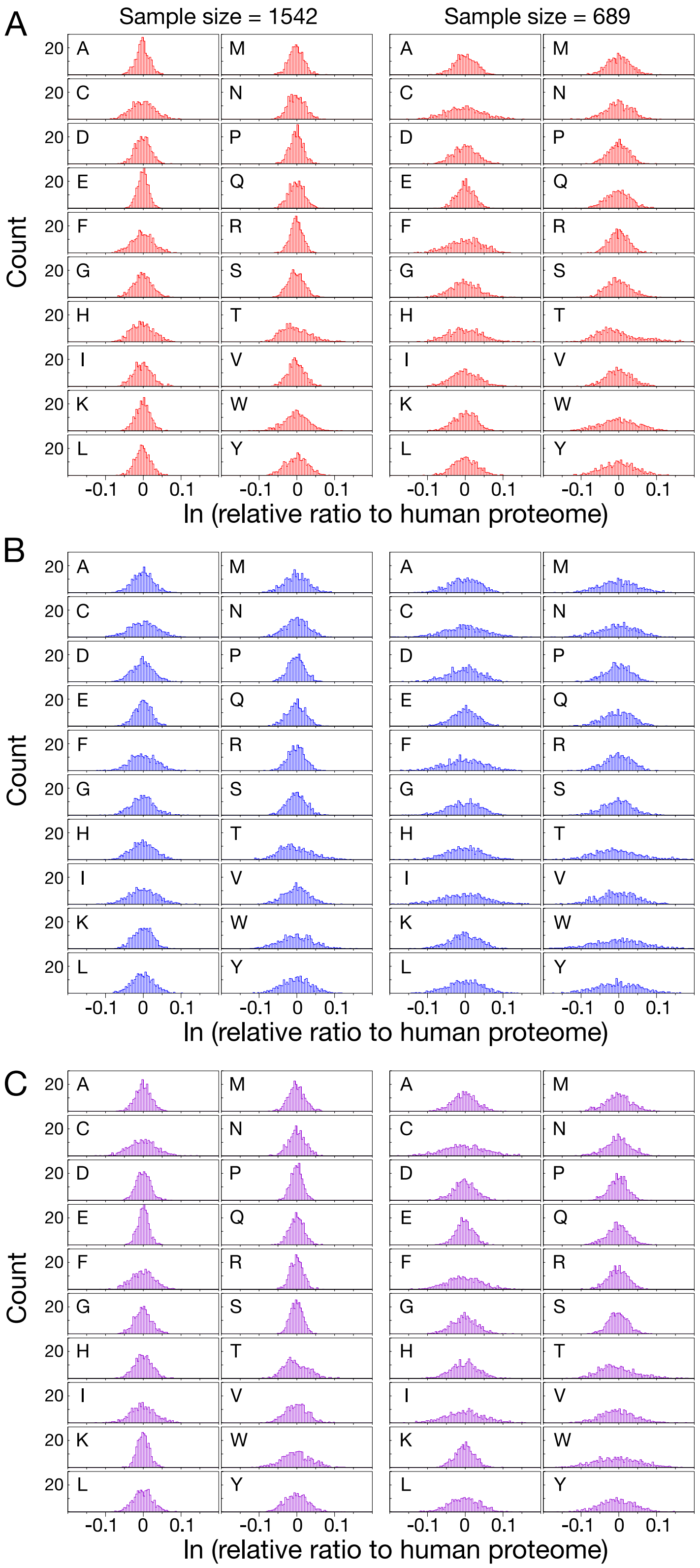


**Figure S1.** Distribution of the log-odds ratio relative to the human proteome of being in an intrinsically disordered region as predicted by the (A) VLXT, (B) VL3, and (C) VSL2 algorithm for randomly selected subgroups of the human proteome (left, N =1542; right, N = 689)

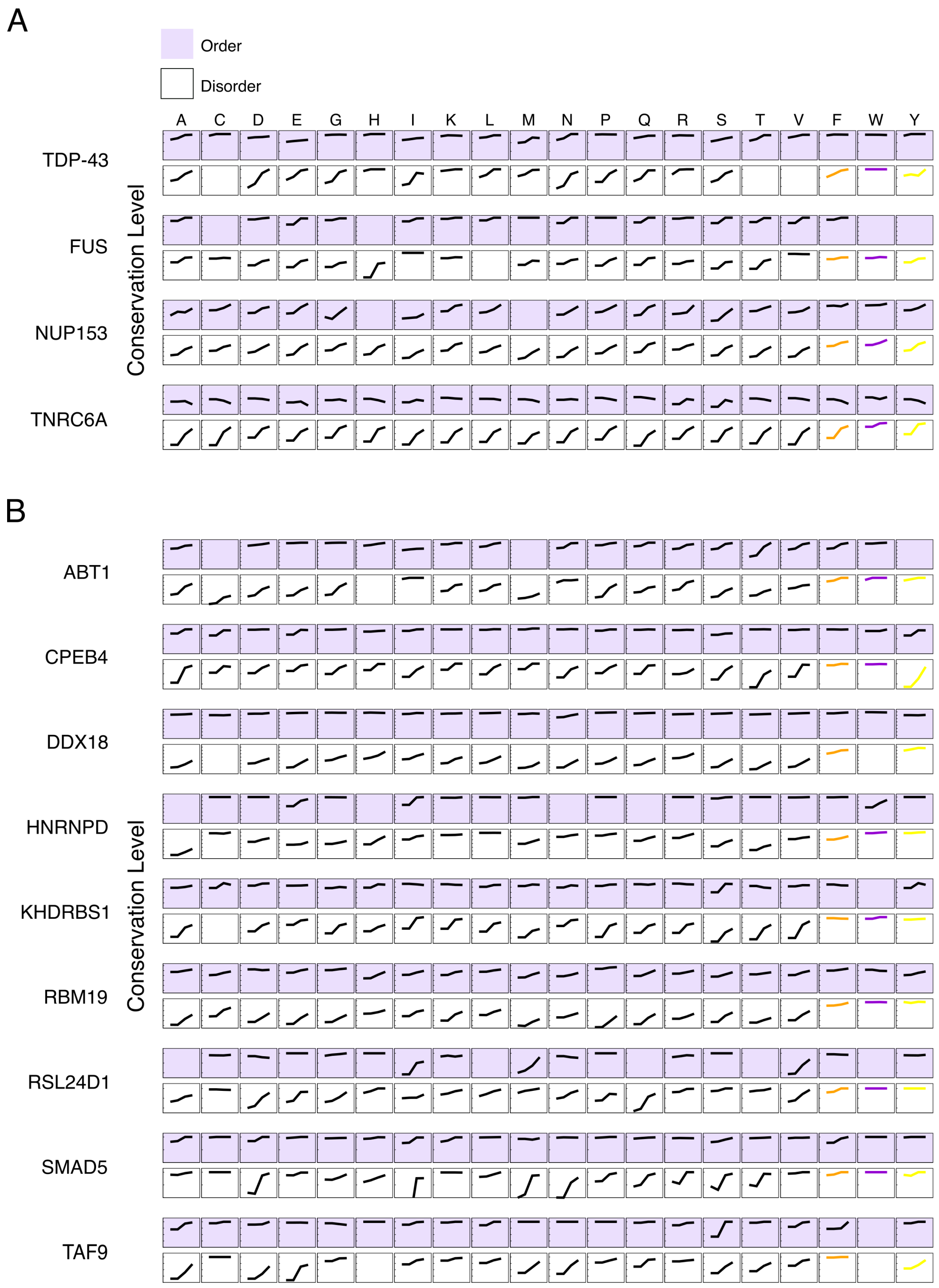


**Figure S2.** Average level of sequence conservation by amino acid type for residues in ordered (purple) or disordered regions (white) as a function of decreasing taxonomic ranks (Chordata, Vertebrata, Tetrapoda, Mammalia), for (A) the example proteins presented in fig. 2 of the main text and (B) the example proteins presented in figure S3 (below). Only residues in disordered (ordered) regions longer than 40 (20) consecutive amino acids were considered.

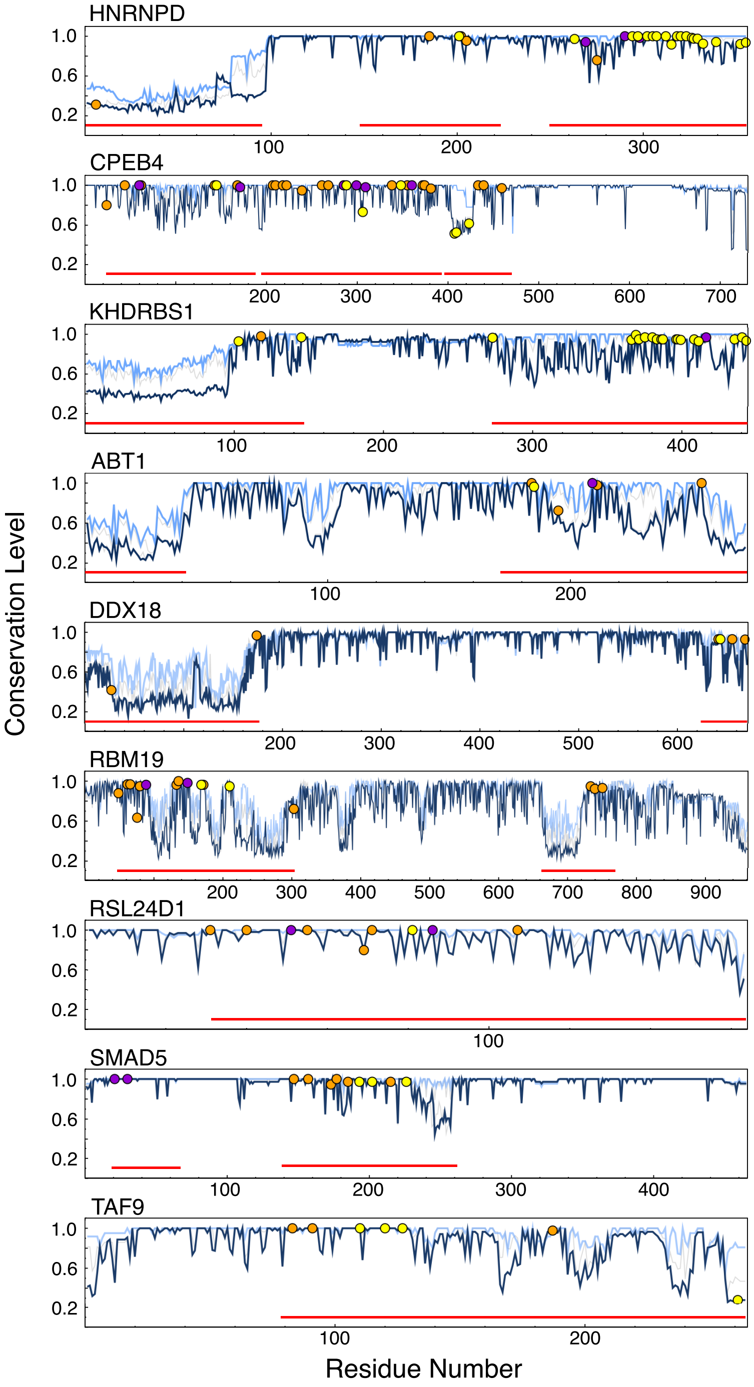


**Figure S3.** Sequence conservation in example proteins with highly conserved aromatic residues. Levels of sequence conservation quantified by the Shannon entropy: a value of 1 means that all the residues aligned at a given position are identical. Levels of conservation in chordates (dark blue), vertebrates (gray), tetrapods (gray), and mammals (light blue) are plotted versus the corresponding residue number in the human sequence. Predicted disordered regions longer than 40 residues are indicated with red bars. Aromatic residues are labeled on the chordate line: Phe (orange), Trp (purple), Tyr (yellow).
